# Non-invasive vagus nerve stimulation decreases vagally mediated heart rate variability

**DOI:** 10.1101/2023.05.30.542695

**Authors:** Kristin Kaduk, Alessandro Petrella, Sophie J. Müller, Julian Koenig, Nils B. Kroemer

## Abstract

The vagus nerve plays a critical role in balancing the body’s physiological functions, including the cardiovascular system. Measures of heart rate (HR) and its variability (HRV) may provide non-invasive proxies of vagal activity in humans, but transcutaneous auricular vagus nerve stimulation (taVNS) has produced mixed effects so far—limited by a lack of studies stimulating the right branch. Here, we used a randomized cross-over design to study the effects of taVNS on HR and HRV. To estimate how the side of the stimulation (left vs. right ear) affects cardiovascular function, we recorded an electrocardiogram in four sessions per person (factors: Stimulation × Side). To evaluate potential interactions with physiological states, we investigated three phases per session: baseline, during stimulation (taVNS vs. sham), and after consuming a milkshake (∼400 kcal) with concurrent stimulation. First, we found moderate evidence against an effect of taVNS on HR (BF_10_=0.21). Second, taVNS decreased HRV (multivariate *p* =.004) independent of physiological state with strong evidence for RMSSD (BF_10_=15.11) and HF-HRV (BF_10_=11.80). Third, taVNS-induced changes were comparable across sides and more strongly correlated (vs. sham), indicating similar cardiovascular effects independent of the stimulation side. We conclude that taVNS reduces HRV without altering HR, contradicting the common assumption that increased HRV indexes a heightened vagal tone. Instead, our results support a putative role of vagal afferent activation in arousal. Crucially, modulatory effects on the cardiovascular system can be safely elicited by taVNS on both sides, opening new options for treatment.

**Graphical Abstract:** Created with BioRender.com

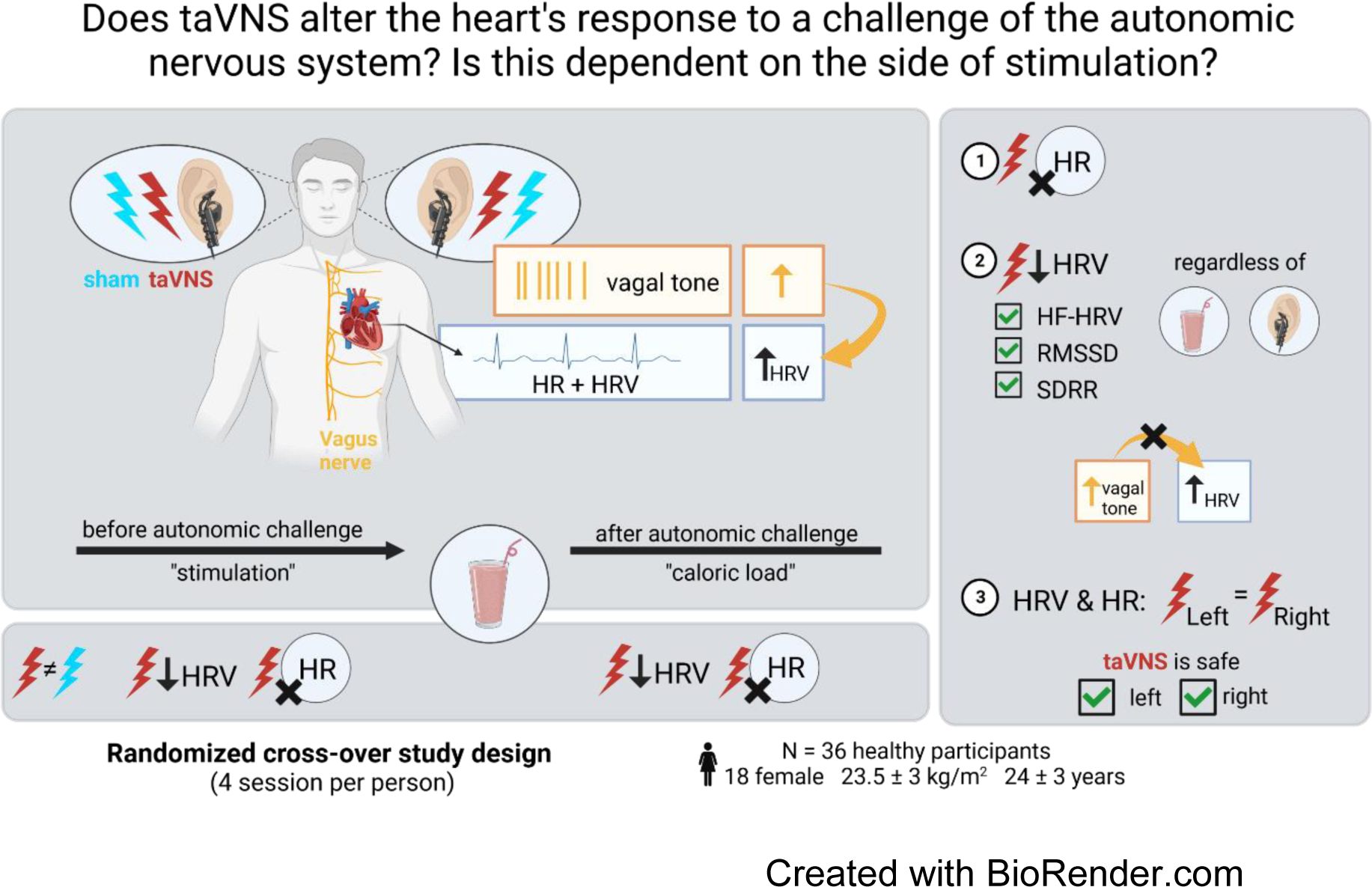

## Introduction

The communication between the brain and the peripheral organs, such as the heart, plays a crucial role in maintaining the body’s physiological and metabolic homeostasis (Capilupi et al., 2020). The peripheral organs send interoceptive signals to the brainstem’s nucleus of the solitary tract (NTS) through vagal afferents to convey the body’s states, such as hunger or alertness (Havel, 2001; Maniscalco & Rinaman, 2018). Brain signals are then transmitted to peripheral organs via vagal efferents influencing the gastric frequency (Hong et al., 2019; Teckentrup et al., 2020) or the outflow of the heart by adjusting the firing rate of the pacemaker: the sinus node (Allen et al., 2022; Gourine et al., 2016; Petzschner et al., 2021). Consequently, animal studies have shown that (invasive) stimulation of both right and left cervical vagus nerves affects heart rate (HR) and heart rate variability (HRV), with greater effects often seen on the right side (Huang et al., 2010; Lee et al., 2018; Yoshida et al., 2018).

Despite the vital relevance of brain–body interactions for adaptive human behavior, the impact of vagus nerve stimulation (VNS) on physiological processes and its role in ensuring energy homeostasis is not fully understood <u>(see Burger et al., 2020;</u> <u>Wolf et al., 2021)</u>. For example, transcutaneous auricular vagus nerve stimulation (taVNS) allows studying the interaction between the vagus nerve and the heart in humans by non-invasively stimulating the auricular branch of the vagus nerve in the ear (Butt et al., 2020; Farmer et al., 2021). These acute taVNS studies have yielded mixed results on HRV, with some studies finding increased HRV from stimulating the left side (Antonino et al., 2017; Forte et al., 2022; Geng et al., 2022) or the right side (De Couck et al., 2017; Gauthey et al., 2020; Machetanz et al., 2021). In contrast, other studies observed a decrease in HRV (Altınkaya et al., 2023; for LF/HF ratio: Clancy et al., 2014; Weise et al., 2015), whereas most studies found no effect on HRV (e.g., Borges et al., 2019; Burger et al., 2019; Galli et al., 2003; Jansen et al., 2011; Šinkovec et al., 2023). This heterogeneity is reflected in a recent meta-analysis, concluding that taVNS does not robustly alter vagally-mediated HRV in humans (Wolf et al., 2021).

Notwithstanding the inconsistent results across studies, there are open questions concerning the side of stimulation and the body’s metabolic state during stimulation. Most taVNS studies have stimulated the left ear, and only one study with right-sided stimulation and two studies with bilateral stimulation (Bretherton et al., 2019; Clancy et al., 2014; De Couck et al., 2017) could be included in the meta-analysis (Wolf et al., 2021). Fewer studies use right-sided taVNS due to a hypothesized higher risk of cardiovascular side effects, such as bradycardia (Kim et al., 2022) and the predominant innervation of the sinus-atrial node by the right vagus nerve (Ardell & Randall, 1986). However, the signal from both auricular branches of the vagus nerve is integrated before activating vagal efferents to the heart, suggesting that side effects may be negligible (M. Chen et al., 2015; Kim et al., 2022; Redgrave et al., 2018). In addition, small sample sizes <u>(Wolf et al., 2021,</u> meta-analysis: median(N) = 30, range(N) = 7-60 of taVNS studies with within-subject design), lack of a common stimulation protocol, appropriate baseline measurements, and adequate control conditions may contribute to discrepancies across studies. Likewise, other physiological factors, such as the respiratory rate or hormonal balance (Kozorosky et al., 2022; Sclocco et al., 2019; Szulczewski, 2022), interact with the effect of taVNS, indicating a complex interplay between physiological states and taVNS-induced changes. To summarize, pressing questions about the differential effects of taVNS at the left vs. the right ear and the emerging evidence of interactions with metabolic state (Altınkaya et al., 2023; Vosseler et al., 2020) call for additional studies on cardiovascular effects. To close the gap, we employed a randomized cross-over design to investigate the effect of stimulation (taVNS vs. sham) on HR and HRV on both auricular branches of the vagus nerve (left vs. right) in different metabolic states, both before and after consuming a milkshake (∼400 kcal).

## Methods

### Participants

The study was preregistered on the Open Science Framework (https://doi.org/10.17605/OSF.IO/26V5N). The sample size was selected to provide at least a power of 1-β = .90 for small-to-medium-sized within-subject effects (Cohen’s *f* = .15, *dz* ∼.50), leading to a lower-bound estimate of 34 participants after quality control. A total of 38 participants were invited to participate. One participant was excluded due to paraesthesia experienced in the sham condition, and another participant was excluded due to undetectable R-peaks in two sessions. Thus, 36 participants were included in the final analysis (18 women, M_age_ = 24 ± 3 years, BMI: 23.5 ± 2.6 kg/m^2^). According to screenings, all participants were physically healthy and excluded if they had contraindications for taVNS (e.g., irremovable earrings or piercings on the ear). After completing the fourth session, participants received a fixed compensation of 80€ or partial course credits of 4 h and 40€. All participants provided their written informed consent before the experiment. The ethics committee of the Faculty of Medicine at the University of Tübingen approved the experiment, and all procedures were carried out in accordance with the Declaration of Helsinki.

### Procedure

To maximize power and account for inter-individual variability, we adopted a randomized cross-over design. The three within-subject factors were stimulation side (right, left), stimulation (taVNS, sham), and time (baseline, stimulation, caloric load), leading to four sessions per participant. Each session had the same timeline (Fig. 1A), starting with a no-stimulation baseline phase while recording the ECG for 30 min. To collect the ECG, we continuously measured the heart’s electrical activity at 5000 Hz using three bipolar electrodes connected to a BrainAmp amplifier (Brain Products, Germany). Electrodes that shared the same negative derivation were placed on the left and right sides of the clavicle pits. The positive derivations were set on the left side in the seventh intercostal space, as previously described by Teckentrup et al., 2020. ECG recordings continued while participants received taVNS or sham stimulation for 30 min before and after administering a standardized caloric load (∼400 kcal milkshake, details in SI). Participants were instructed to attend the session 2-3h after their last meal. To control for circadian rhythms, the four experimental sessions were scheduled at approximately the same time on a different day. Except for state questions every 15 min, participants were resting and listening to an audiobook.

**Figure 1.**
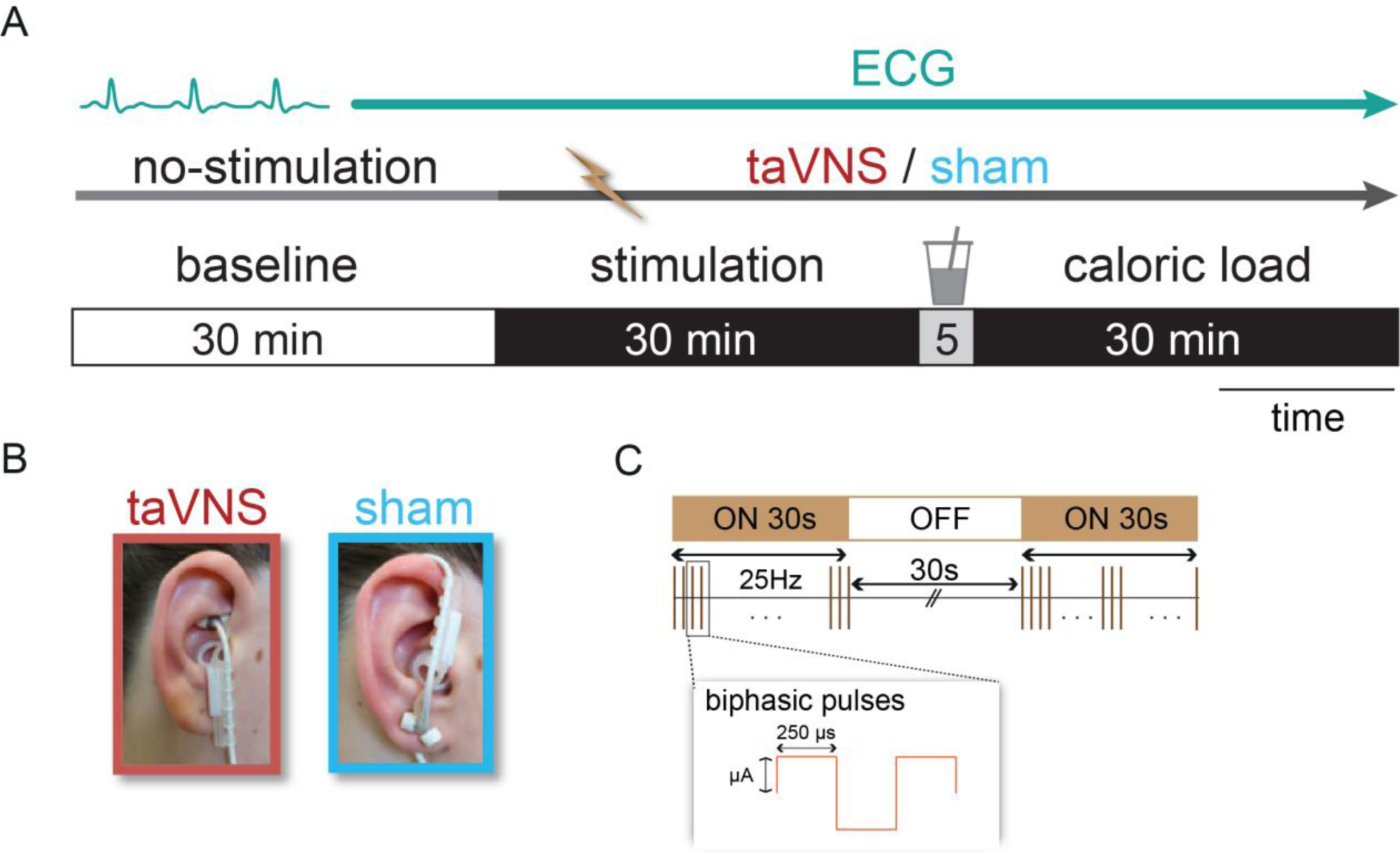
Schematic summary of the experiment. A: The timeline of a session includes the phases of baseline (30 min, no stimulation) followed by stimulation (30 min) when either taVNS or sham was applied. Participants received a high-caloric load to challenge the autonomic system, while we collected the phase after the caloric load (30 min) with concurrent stimulation. B: The electrode locations at the cymba conchae for the transcutaneous auricular vagus nerve stimulation (taVNS, red) and the ear lobe for sham stimulation (blue) are displayed. C: Illustration of the stimulation protocol with a biphasic impulse frequency of 25 Hz, the pulse width of 250 μs with alternating intervals of 30 s stimulation on and 30 s stimulation off. The stimulation intensity (mA) was adjusted for each participant until they felt a “mild pricking”.

### Electrical stimulation

To stimulate the auricular branch of the vagus nerve, we used the NEMOS® device (cerbomed GmbH, Erlangen, Germany) following the procedure of previous studies (Ferstl et al., 2021; Frangos et al., 2015; Neuser et al., 2020; Teckentrup et al., 2021). For taVNS, the electrode was placed at the cymba conchae (Peuker & Filler, 2002) as it induces robust activation in the NTS (Borgmann et al., 2021; Frangos et al., 2015; Teckentrup et al., 2021; Yakunina et al., 2017). The electrode was turned upside down for sham stimulation and placed at the earlobe (Butt et al., 2020) (Fig. 1B). Analogous to previous studies, each participant’s stimulation intensity was adjusted to a “mild pricking” level (VAS = 5) before task onset (see Fig. 1C and Table S1).

The side of stimulation was randomized in advance. To ensure that taVNS and sham are comparable on each side, the stimulation side was not changed between Sessions 1 and 2 (i.e., if a participant received left-sided taVNS in Session 1, they received sham on the left side in Session 2). Participants were blinded regarding the stimulation condition and after each session, they guessed which condition they had received.

### Data analysis

### R-peak detection & cardiovascular indices

All cardiovascular indices were derived from continuous ECG recordings. The ECG signal was extracted and down-sampled to 1000 Hz using the FieldTrip toolbox (Oostenveld et al., 2011, http://fieldtriptoolbox.org). Custom code (https://github.com/dagdpz/body_signals_analysis) was partly adjusted and used for the pre-processing and R-peak detection. We processed the recorded and detrended ECG data to remove the 50 Hz power line interference (19^th^ order Butterworth filter with a passband of 40 Hz, stop band of 100 Hz, passband ripple of 1 dB, and stopband ripple of 150 dB) and baseline drifts (high-pass filter with a cut-off frequency of 0.5 Hz), which can affect the accuracy of peak detection. We then detected the R peaks and QRS complex and computed the R-R interval time series. As the ECG morphology is strongly affected by movement artifacts, we used an automatic procedure to check the R peaks and the R-R intervals for robustness and deviations (see SI). All detected deviations and their consecutive R-R intervals were deleted from the signal. On average, 97.1% (1769s, range: 81% - 100%) of a 30 min block of ECG recordings was further analyzed to derive cardiovascular indices (except for one phase with 956 s of ECG data).

HRV is the variation in time between consecutive heartbeats. Four standard HRV measures in taVNS studies are SDRR, RMSSD, HF-HRV, and LF/HF ratio (Wolf et al., 2021). We computed the following time-domain measures: the standard deviation of RR intervals (SDRR in ms) was calculated as a general trend in the ECG, and the root mean squares of successive differences of adjacent heartbeats (RMSSD in ms), representing the beat-to-beat variance in the heart period. For frequency-domain HRV, we computed the spectra using the HRV toolbox of the PhysioNet Cardiovascular Signal Toolbox (Vest et al., 2018) in 6-minute windows of 30 s steps, after cubic spline interpolation to account for potential movement artifacts during questionnaires, using Welch’s method. For each time window 20% of data could be rejected before a window is considered too low quality for analysis. Two frequency-domain HRV measurements were computed from R-R intervals based on power spectral density: High-frequency HRV (HF-HRV) at 0.15-0.4 Hz and low-frequency HRV (LF HRV) at 0.04-0.15 Hz, following recommendations from the Task Force of the European Society of Cardiology and the North American Society of Pacing and Electrophysiology.

### Statistical analysis

The statistical analysis was performed using R (version 4.1.2, R Core Team, 2022). Cardiovascular indices (HR, HRV) were baseline-corrected by subtracting the individual session-specific baseline index from later indices (separately for stimulation and caloric load phases). To assess the impact of the three within-subject factors (Stimulation (taVNS, sham) × Side (right, left) × Phase (stimulation, caloric load)) on the four dependent HRV indices, we conducted a multivariate analysis of variance (MANOVA) first. Next, we investigated the effects on each HRV index separately. We calculated the net effect of stimulation by subtracting baseline-corrected sham from baseline-corrected taVNS (pairwise differences) for each participant and session. To avoid distributional assumptions of parametric statistics, we bootstrapped their distribution for statistics <u>(e.g. Teckentrup et al., 2020, 50,000 resampling steps)</u>, but mixed-effects models would have led to comparable conclusions. For main effects across sides, individual estimates were averaged first. As a threshold, we used *p* ≤ 0.05 (two-tailed). To quantify the effect of taVNS on HR and HRV, we calculated effect sizes (Cohen’s dz) and carried out Bayesian one-sample t-tests including individual estimates of taVNS-induced changes in HRV per stimulation side (left and right). To reflect that small-to-moderate effects are likely, we used a Cauchy prior with a width of 0.5 in JASP v.0.17.2 (Quintana & Williams, 2018).

## Results

### Stimulation decreases HR, while caloric load increases it

To evaluate the effect of stimulation (taVNS vs. sham; left and right side) before administering the caloric load, we analyzed changes in HR during the 30 min stimulation phase versus baseline. Across both stimulation conditions, HR decreased after stimulation (Fig. 2A; b = 0.82 bpm, 95% CI [−1.35, −0.30], *p_Boot_* = .002). However, we did not observe differences between taVNS and sham across both sides (b = −0.11 bpm, 95% CI [−0.70, 0.46], *p*_Boot_ = .722) before the challenge.

**Figure 2.**
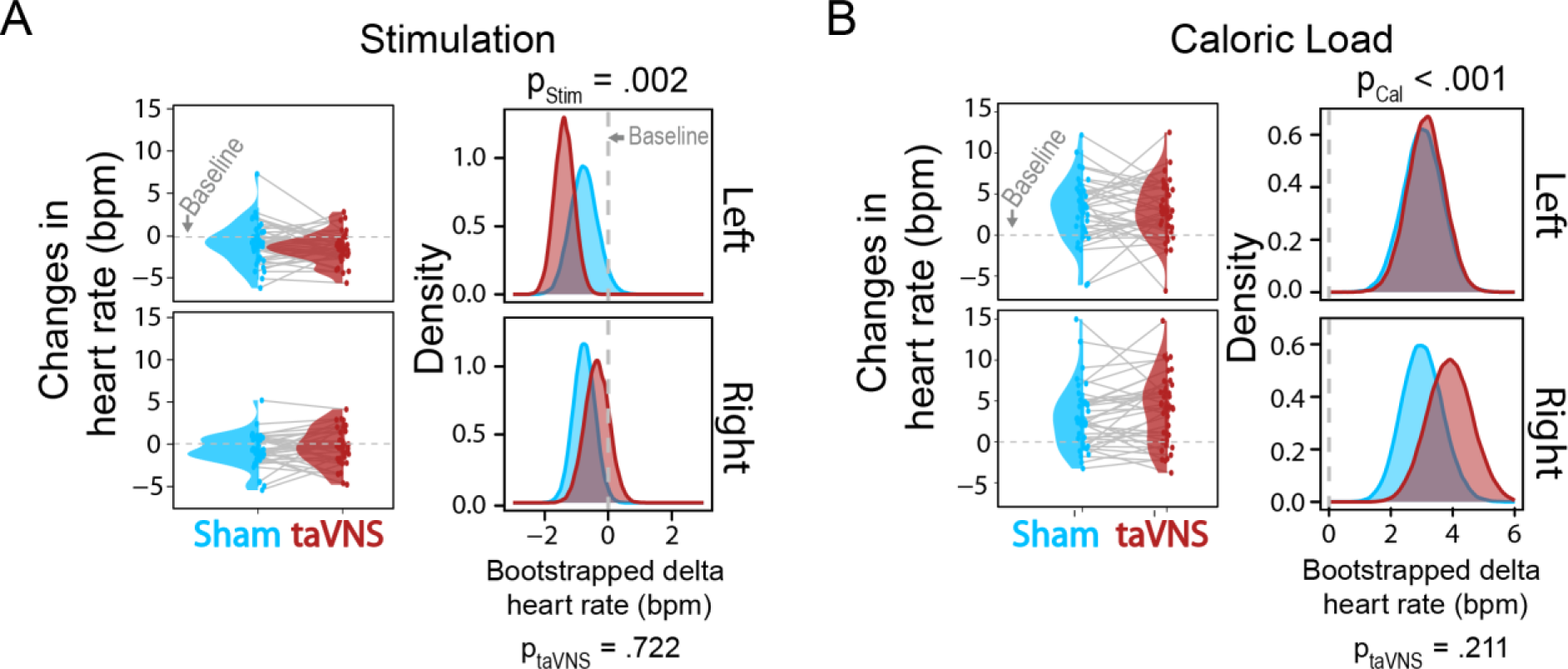
Stimulation decreases and caloric load increases heart rate (HR). (A) During stimulation, HR decreases across both sides (left vs. right) and both stimulation conditions (taVNS in red vs. sham in blue), indicated by the *p*-values below the heading. We observed no differences between stimulation conditions on each side, indicated by the bootstrapped distributions of changes relative to baseline. (B) After the caloric load, HR increases across both sides (left and right) and stimulation conditions (taVNS and sham), as indicated by the *p*-values below the heading. We observed no differences between stimulation conditions on each side, indicated by the bootstrapped distributions of changes relative to baseline. The dots depict each participant’s change in average HR for taVNS (in red) and sham (in blue) compared to the baseline.

Next, to evaluate the interaction with a challenge of the autonomic system, we analyzed the main effect of caloric load on HR. As expected, HR increased after consuming a 400-kcal milkshake (Fig. 2B; b = 3.28 bpm, 95% CI [2.33, 4.21], *p_Boot_* <.001; relative to the stimulation phase: b = 4.09 bpm, 95% CI [3.36, 4.85], *p_Boot_* <.001). We also did not observe differences between taVNS and sham across both sides (b = 0.49 bpm, 95% CI [−0.28, 1.25], *p_Boot_* = .211). These results suggest that stimulation and caloric load altered HR within a session, but these changes were not specific to taVNS, as suggested by the moderate evidence against an effect (Bayes factor, BF_10_ = 0.21).

### taVNS reduces HRV before a challenge

To compare taVNS-induced effects on HRV with prior studies, we computed four commonly reported HRV measures used in most taVNS studies (Wolf et al., 2021). Since HRV indices are correlated measures of a similar construct, we initially ran a multivariate analysis to evaluate taVNS-induced changes among all HRV indices. HRV was modulated by both the stimulation (MANOVA, *V*_Pillai’s Trace_ = 0.053, *F*(4,276) = 3.88, *p* = .004) and metabolic state (*V*_Pillai’s Trace_ = 0.215, *F*(4,277) = 18.97, *p* < .001), but the side of stimulation did not contribute to the changes in HRV and there was no interaction of stimulation with metabolic state (*p* > .05).

To complement these multivariate findings, we investigated the effect of taVNS on the individual HRV indices by phase (i.e., analogous to post hoc tests). Before the caloric load, stimulation (i.e., taVNS and sham) increased SDRR (Fig. 3A; b = 5.47 ms, 95% CI [2.90, 8.22], *p_Boot_* < .001), RMSSD (b = 3.68ms, 95% CI [1.40, 6.20], *p_Boot_* = .001) and LF/HF ratio (b = 0.13, 95% CI [0.05, 0.22], *p_Boot_* = .001), but not HF-HRV (b = 38.83 ms^2^, 95% CI [−63.21, 140.89], *p_Boot_* = .460).

**Figure 3.**
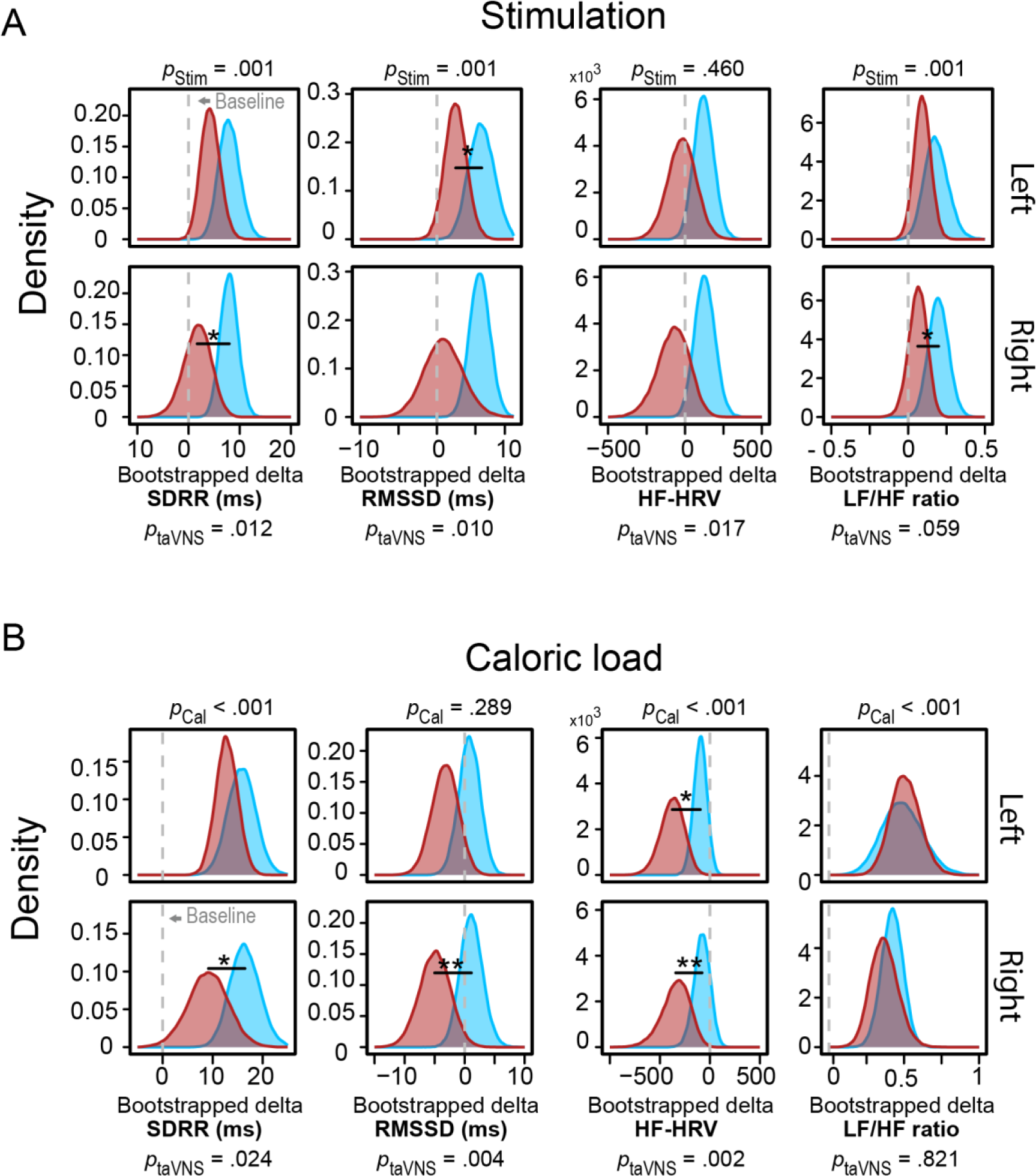
taVNS decreases HRV. A: Across both stimulation conditions (taVNS vs. sham), LF/HF ratio, SDRR and RMSSD are altered across both sides (left vs. right) and both stimulation conditions (taVNS in red vs. sham in blue), indicated by the *p*-values above each plot. taVNS (red distribution) decreased HRV compared to sham (blue distribution) before the challenge, indicated by the bootstrapped distributions of the delta to the baseline separated for each side. Statistics for the taVNS-induced effects across both sides are displayed below the panel. B: The caloric load alters HRV, except for RMSSD, across both sides (left vs. right) and stimulation conditions (taVNS vs. sham), indicated by the *p*-values above each plot. taVNS decreased SDRR, RMSSD and HF-HRV after the caloric load, as indicated by the bootstrapped distributions.

Notably, in the time domain, we observed a taVNS-induced decrease in SDRR (Fig. 3A; b = −4.98 ms, 95% CI 9.56, −1.04], *p_Boot_* = .012) and RMSSD across both sides (b = −4.05 ms, 95% CI [−7.44, −0.92], *p_Boot_* = .010). In the frequency domain, taVNS reduced HF-HRV across sides (b = −174.84 ms^2^, 95% CI [−365.75, −24.65], *p_Boot_* = .017), which was more pronounced at the right ear (Table 1, see Fig. S1). To summarize, HRV decreases during taVNS compared to sham and there was no interaction with stimulation side.

**Table 1.** Comparison taVNS vs. sham during stimulation phase after bootstrapping.

| Index | Both sides |  |  | Left |  | Right |  |
| --- | --- | --- | --- | --- | --- | --- | --- |
|  | mean<br>[CI<sub>L</sub>, CI<sub>U</sub>] | *p* | effect<br>size *d*<sub>z</sub> | mean<br>[CI<sub>L</sub>, CI<sub>U</sub>] | *p* | mean<br>[CI<sub>L</sub>, CI<sub>U</sub>] | *p* |
| **Stimulation** |  |  |  |  |  |  |  |
| HR | -0.11<br>[-0.70, 0.46] | .722 | 0.058 | -0.61<br>[-1.40, 0.18] | .128 | 0.40<br>[-0.25, 1.17] | .305 |
| SDRR | **-4.98**<br>[-9.56, -1.04] | **.012** | **0.471** | -3.69<br>[-8.03, 0.60] | .092 | **-6.28**<br>[-12.84, -0.78] | **.022** |
| RMSSD | **-4.05**<br>[-7.44, -0.92] | **.010** | **0.441** | **-3.39**<br>[-6.54, -0.56] | **.017** | -4.70<br>[-10.32, 0.64] | .088 |
| HF- HRV | **-174.84**<br>[-365.75, -24.65] | **.017** | **0.415** | -143.75<br>[-353.64, 45.97] | .147 | -206.04<br>[-448.15, 2.88] | .054 |
| LF/HF<br>ratio | -0.11<br>[-0.25, 0.00] | .059 | 0.347 | -0.09<br>[-0.27, 0.08] | .300 | **-0.14**<br>[-0.29, -0.00] | **.048** |
| **Caloric Load** |  |  |  |  |  |  |  |
| HR | 0.49<br>[-0.28, 1.25] | .211 | -0.155 | 0.10<br>[-1.26, 1.42] | .882 | 0.89<br>[-0.24, 1.98] | .123 |
| SDRR | **-5.16**<br>[-10.07, -0.63] | **.024** | **0.318** | -3.02<br>[-7.48, 1.44] | .184 | **-7.25**<br>[-14.91, -0.51] | **.034** |
| RMSSD | **-5.09**<br>[-8.83, -1.54] | **.004** | **0.460** | -4.07<br>[-9.02, 0.76] | .101 | **-6.13**<br>[-11.02, -1.63] | **.006** |
| HF- HRV | **-271.54**<br>[-477.26, -95.60] | **.002** | **0.447** | **-279.61**<br>[-550.76, -30.18] | **.027** | **-264.05**<br>[-475.41, -75.68] | **.005** |
| LF/HF<br>ratio | -0.02<br>[-0.16, 0.13] | .821 | 0.032 | 0.02<br>[-0.19, 0.23] | .845 | -0.05<br>[-0.20, 0.10] | .492 |

Note. CI<sub>L</sub> = 95% lower bound and CI<sub>U</sub> = 95% upper bound of confidence interval

### taVNS robustly affected SDRR, RMSSD and HF-HRV after a caloric load

To investigate HRV after an autonomic challenge, we first evaluated the effect of a standardized caloric load on HRV. After the challenge, we observed a decrease in HF-HRV (Fig. 3B; b = −223.29 ms^2^, 95% CI [−378.31, −92.20], *p_Boot_* < .001) and an increase in SDRR (b = 13.56 ms, 95% CI [9.02, 18.29], *p_Boot_* < .001), and LF/HF ratio (b = 0.45, 95% CI [0.31, 0.60], *p_Boot_* < .001). In contrast, the challenge did not affect RMSSD (b = −1.58 ms, 95% CI [−4.63, 1.26], *p_Boot_* = .289).

Regarding the time-domain measures of HRV, taVNS reduced SDRR (Fig. 3B, b = −5.16 ms, 95% CI [−10.07, −0.63], *p_Boot_* = .024) and RMSSD across both sides (b = −5.09 ms, [−8.83, −1.54], *p_Boot_* = .004) after the caloric load. Regarding the frequency-domain measures of HRV, taVNS reduced HF-HRV after the caloric load across both sides (b = −271.54 ms^2^, [−477.26, −95.60], *p_Boot_* = .002) and also for each side (Table 1). However, this did not lead to taVNS-induced changes in LF/HF ratio across both sides (b = −0.02, [−0.16, 0.13], *p_Boot_* = .821). To summarize, taVNS at both ears reduced SDRR, RMSSD, and HF-HRV after the caloric load, replicating the effects in independent sessions within the same sample.

To investigate the potential variance in taVNS effects on HRV across different physiological conditions, we compared HRV between the stimulation and caloric load phase. RMSSD and HF-HRV decreased after the caloric load compared to the stimulation (RMSSD: b = −5.27 ms, 95% CI [−8.00, −2.59], *p_Boot_* < .001; HF-HRV: b = - 262.23 ms^2^, 95% CI [−374.95, −154.94], *p_Boot_* < .001) and SDRR and LF-HF ratio increased (SDRR: b = 8.10 ms, 95% CI [3.65, 12.77], *p_Boot_* < .001; LF-HF ratio: b = 0.31, 95% CI [0.21, 0.42], *p_Boot_* < .001). After the caloric load, we found that taVNS decreased HF-HRV and RMSSD, further amplifying the reduction in RMSSD and HF-HRV induced by the milkshake. Notably, the taVNS-induced effects in the caloric load phase did not significantly differ from the taVNS-induced effects in the stimulation phase for any HRV index (*p_Boot_* > .05, see Table S2). To summarize, taVNS consistently decreased HRV, irrespective of the physiological state, as shown by strong evidence, particularly for RMSSD (BF_10_ = 15.11) and HF-HRV (BF_10_ = 11.80).

### No side-specific differences of HR and HRV due to taVNS

To better understand the lateralization effects of taVNS, we investigated whether left or right taVNS affects HR or HRV differently. In contrast to the theorized larger risk of right taVNS for cardiac function, there was no difference in HRV (Fig. 4A; *p*s *_Boot_* > .05) or HR (Fig. 4B; *p _Boot_* > .05), indicating similar cardiovascular effects across both sides. For 9 out of 10 computed cardiovascular indices, the changes in HRV (Fig. 4C; see Table S3) and HR (Fig. 4D; see Table S3) correlated more strongly across sides for taVNS vs. sham (one-sample t-test of the Fisher z-transformed coefficients, *t*(9) = 3.16, *p* =.012, Fig. 4E).

**Figure 4.**
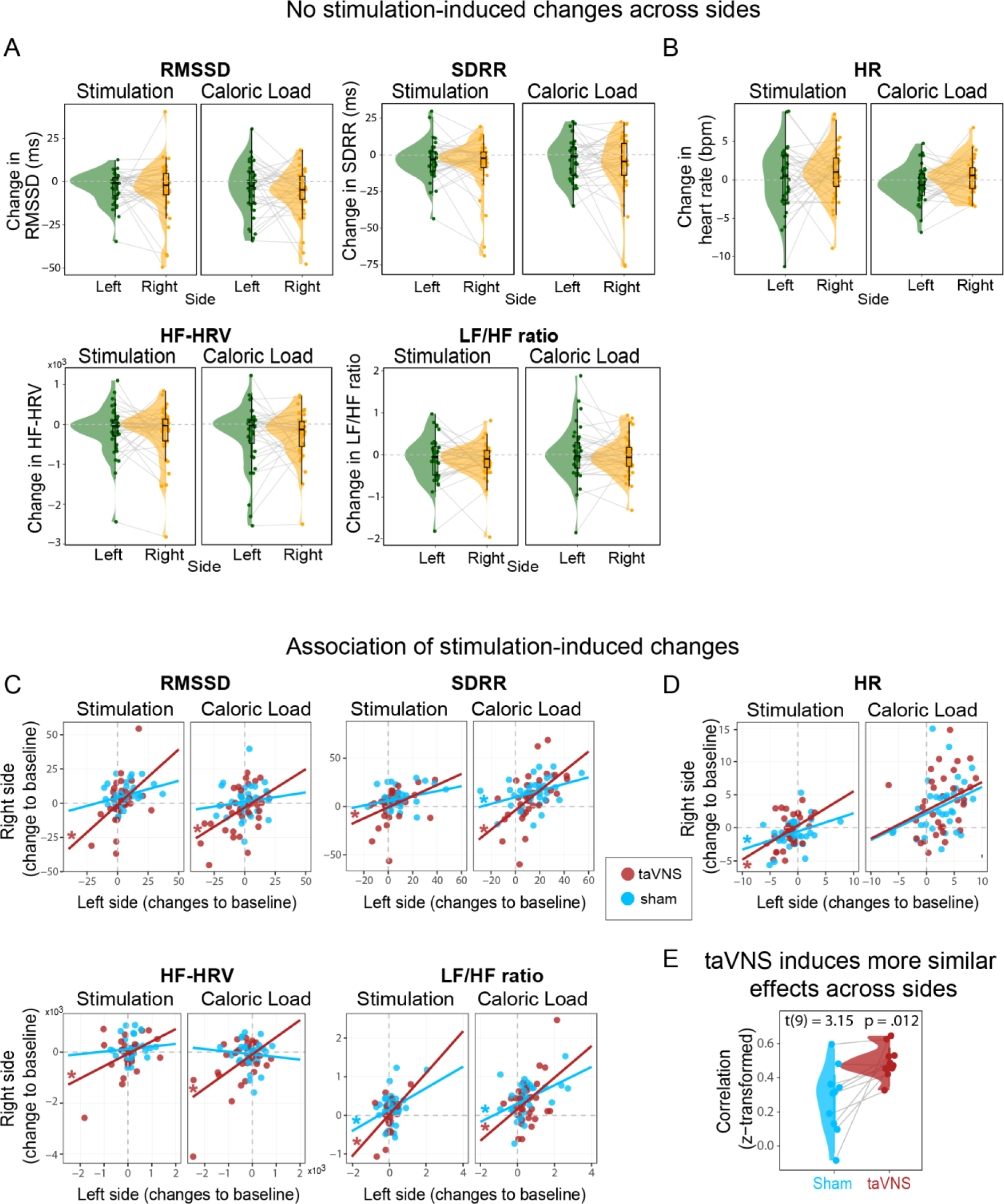
No differences between the sides of vagus nerve stimulation in heart rate (HR) and heart rate variability (HRV). A&B: For cardiovascular indices, HRV and HR depict the participant’s average score (taVNS – sham of their change from baseline), and the group distribution shows no difference for the side-specific comparison (left vs. right) for either the stimulation or caloric load phases. C&D: taVNS-induced changes (red) in HRV and HR are more strongly correlated across sides compared to sham (blue), indicated by the stars at the correlation lines for the data of each participant. E: Higher correlation coefficients for cardiovascular indices between sides of stimulation after taVNS compared to sham.

## Discussion

The cardiovascular system is regulated by bidirectional communication via the vagus nerve with the brain. Although HRV is thought to be closely linked to vagal tone (Laborde et al., 2017; Pomeranz et al., 1985), previous studies investigating the effect of non-invasive taVNS on HRV in humans have yielded inconclusive results. To address this gap, we used a randomized cross-over design to evaluate the side- and state-specific effects of taVNS on changes in HR and HRV (i.e., within participants). Although the stimulation and the caloric load as experimental manipulations altered HR, we found no effect of taVNS (vs. sham). Contrary to theorized associations of heightened vagal tone with greater HRV, stimulating vagal afferents did not increase HRV. Instead, taVNS decreased SDRR, RMSSD and HF-HRV before and after the caloric load, indicating a successful modulation of cardiac function in both states. Intriguingly, we observed comparable taVNS-induced changes across sides that were more strongly correlated than sham-induced changes. This supports the interpretation that side-specific vagal afferent inputs through the auricular branch are integrated into the brain for cardiovascular control before relaying them efferently to the body. We conclude that taVNS-induced changes in HR and HRV are largely independent of the side of the stimulation and the body’s metabolic state. Crucially, our results may help reconcile acute modulatory effects of the cardiovascular system facilitating arousal, not relaxation, with VNS-induced increases in pupil dilation (D’Agostini et al., 2023; Lloyd et al., 2023; Sharon et al., 2021), invigoration (Neuser et al., 2020) and VNS-induced decreases in alpha oscillations in an EEG (Y. Chen et al., 2023; Lewine et al., 2019; Sharon et al., 2021), that are not well explained by parasympathetic activation of a “rest and digest” mode.

Consistent with recent findings, we observed a robust taVNS-induced reduction in SDRR, HF-HRV and RMSSD across both sides (Altınkaya et al., 2023; Bretherton et al., 2019; Clancy et al., 2014; Weise et al., 2015). These findings contradict the conventional assumption that higher HRV indexes total vagal activity or vagal tone (Buchman et al., 2002; Shaffer & Ginsberg, 2017). Notably, this assumption was challenged by recent invasive vagus nerve recordings demonstrating that direct measures of vagal tone are not associated with HRV in rodents (Marmerstein et al., 2021). Relatedly, it has been proposed that HRV is associated with signals from cardiac vagal efferents and that, specifically, HF-HRV reflects the cardiac parasympathetic modulations of vagal efferents and baroreceptor afferent signaling (Eckberg, 1983; Grossman & Taylor, 2007; Hedman et al., 1995). Whereas tonic decreases in HRV may reflect impairments in the body’s autonomic function (Coopmans et al., 2020; Laborde et al., 2017; Thayer & Lane, 2007), an acute decrease in HRV can be adaptive, reflecting an individual’s capacity to allocate energy to cope with demand (Dickerson & Kemeny, 2004; Thayer et al., 2009). Intriguingly, such demand may correspond with vagal afferent signals affecting arousal and motivational drive (Y. Chen et al., 2023; D’Agostini et al., 2023; Neuser et al., 2020; Rong et al., 2014). Notably, previous studies on taVNS-induced changes in HRV yielded mixed results leading to non-significant taVNS-induced changes across published studies so far (Wolf et al., 2021). It is conceivable that differences in study design and taVNS protocols contribute to heterogeneity across studies and it is important to identify potential moderator and mediator variables. Nevertheless, we find consistent effects across complementary indices of HRV, physiological states, and sides of the stimulation, effectively providing a replication of taVNS-induced changes from independent sessions within the study.

An important factor that might modulate the effectiveness of taVNS in regulating cardiovascular function is the body’s metabolic state (Kozorosky et al., 2022; Sclocco et al., 2019; Szulczewski, 2022). After the caloric load, we found that taVNS decreased HF-HRV and RMSSD further amplifying the reduction in RMSSD and HF-HRV induced by the milkshake. Notably, while SDRR and LF/HF ratio increased after the caloric load, taVNS still lead to a reduction in SDRR, with no effect on LF/HF ratio. Likewise, other studies have reported either a decrease in HF-HRV after a caloric load (Lu et al., 1999; Ohara et al., 2015) or an increase in LF/HF ratio after an oral glucose tolerance test (Vosseler et al., 2020). Early stages of digestion may be linked to vagal withdrawal and reduced parasympathetic activity inhibiting the heart’s pacemaker (Ng et al., 2001), resulting in an increased HR. Vagally-mediated processes are crucial to calibrate energy metabolism after increases in glucose levels to regulate eating behavior according to metabolic demands (Berthoud, 2008; Berthoud et al., 2021; Prescott & Liberles, 2022). Of note, two previous studies reported either non-significant or inconclusive taVNS-induced changes in HRV after administering an oral glucose tolerance test (∼300 kcal) (Vosseler et al., 2020) or a milkshake (Altınkaya et al., 2023) using a comparable stimulation protocol. Since these studies were considerably smaller (*N*s = 14 and 15), these differences could be mostly due to power (i.e., increasing N to 36 improves power for moderate within-subject effects dz∼.6 from 58% to 94%). We conclude that taVNS decreased HRV before and after the caloric load even though the load had opposing effects on HRV indices, indicating that taVNS-induced changes are largely independent of metabolic state.

Crucially, our study also provides much-needed information regarding the debate about side-specific effects of tVNS (Kim et al., 2022) on cardiovascular function with important implications for its safety. In contrast to concerns about potential increases in the risk for bradycardia due to parasympathetic hyperstimulation (e.g., due to a case report for long-term VNS stimulation: Pascual, 2015; Shankar et al., 2013), we only found support for acutely induced reductions in HRV, not HR. In general, reports of adverse cardiac effects in healthy young individuals after right taVNS are scarce and do not support a clinically relevant side-specific modulation (Redgrave et al., 2018). By capitalizing on our within-subject design, we quantitatively examined the hypothesized lateralization of taVNS-induced changes in HR and HRV. We showed that taVNS-induced changes in HR and HRV are comparable across sides, providing conclusive evidence against large side-specific effects. While previous studies have reported an increase in HRV with right cymba taVNS as compared to the left side (De Couck et al., 2017; Machetanz et al., 2021), the former used no stimulation as reference (De Couck et al., 2017) and the latter used cavum conchae sham in a between-subject design (Machetanz et al., 2021). Moreover, we found that taVNS-induced changes in HR and HRV between sides were more similar compared to sham-induced changes, indicating that they likely recruited a comparable efferent pathway. Taken together, our findings demonstrate that taVNS elicits modulatory effects on the cardiovascular system that are comparable on both sides, suggesting that right taVNS can be used as safely as left taVNS.

Despite producing converging results across physiological states and stimulation sides, our taVNS study has limitations. First, we used the conventional continuous stimulation protocol and pulsed taVNS (Keute et al., 2021), or low-intensity settings (Šinkovec et al., 2023) may exert different effects on cardiovascular function. Second, future studies may extend our work by recording additional physiological parameters, such as respiration, as HF-HRV and breathing are closely linked (Grossman & Taylor, 2007; Jensen et al., 2022; Soer et al., 2021). Third, future studies could benefit from incorporating additional biomarkers, such as salivary alpha-amylase (Bach, 2014; Giraudier et al., 2022), or muscle sympathetic nerve activity (Clancy et al., 2014) that provide complementary information on the activity of the sympathetic or parasympathetic system. Such biomarkers may support the interpretation of taVNS-induced effects on HRV (see Clancy et al., 2014; Koenig et al., 2021), thereby advancing our understanding of body–brain regulation (Bates et al., 2023).

To conclude, the vagus nerve plays a crucial role in regulating the cardiovascular system according to demand. Here, we investigated the effects of taVNS on HR and HRV in a randomized cross-over design, considering potential interactions with the side of the stimulation and physiological states (i.e., before and after consuming a milkshake). Our results revealed no taVNS-induced changes in HR, while HRV decreased during stimulation and after the caloric load. These taVNS-induced decreases in HRV contradict the popular theory that a heightened vagal tone after afferent stimulation of the vagus nerve would lead to greater HRV, suggesting that cardiovascular effects might be best contextualized with VNS-induced increases in arousal and motivational drive, instead of relaxation. Moreover, we conclude that taVNS-induced effects on the cardiovascular system can be elicited on both sides and in different metabolic states with similar effectiveness without reducing HR or increasing the risk of brachycardia in heathy participants. Consequently, future applications targeting potentially lateralized modulatory effects of taVNS can likely stimulate the right auricular branch of the vagus nerve as safely as the left branch.

## Acknowledgment

We thank Nora Gerth, Nina Röhm, and Vincent Koepp for help with data acquisition and Vanessa Teckentrup for support in setting up the experiment. The study was supported by the Daimler and Benz Foundation, postdoctoral scholarship 32-04/19 and the Deutsche Forschungsgemeinschaft, DFG KR-4555/7-1 and KR-4555/9-1.

## Author contributions

NBK was responsible for the study concept and design. SM and AP preregistered the study, collected the data, and wrote a draft of the procedures. KK processed the ECG data and performed the data analysis and NBK contributed to the analysis. KK and NBK wrote the manuscript. All authors contributed to the interpretation of findings, provided critical revision of the manuscript for important intellectual content, and approved the final version for publication.

## References

Allen, M., Levy, A., Parr, T., & Friston, K. J. (2022). In the Body’s Eye: The computational anatomy of interoceptive inference. PLOS Computational Biology, 18(9), e1010490. https://doi.org/10.1371/journal.pcbi.1010490

Altınkaya, Z., Öztürk, L., Büyükgüdük, İ., Yanık, H., Yılmaz, D. D., Yar, B., Değirmenci, E., Dal, U., & Veldhuizen, M. G. (2023). Non-invasive vagus nerve stimulation in a hungry state decreases heart rate variability. Physiology & Behavior, 258, 114016. https://doi.org/10.1016/j.physbeh.2022.114016

Antonino, D., Teixeira, A. L., Maia-Lopes, P. M., Souza, M. C., Sabino-Carvalho, J. L., Murray, A. R., Deuchars, J., & Vianna, L. C. (2017). Non-invasive vagus nerve stimulation acutely improves spontaneous cardiac baroreflex sensitivity in healthy young men: A randomized placebo-controlled trial. Brain Stimulation, 10(5), 875–881. https://doi.org/10.1016/j.brs.2017.05.006

Ardell, J. L., & Randall, W. C. (1986). Selective vagal innervation of sinoatrial and atrioventricular nodes in canine heart. American Journal of Physiology-Heart and Circulatory Physiology, 251(4), H764–H773. https://doi.org/10.1152/ajpheart.1986.251.4.H764

Bach, D. R. (2014). Sympathetic nerve activity can be estimated from skin conductance responses—A comment on Henderson et al. (2012). NeuroImage, 84, 122–123. https://doi.org/10.1016/j.neuroimage.2013.08.030

Bates, M. E., Eddie, D., Lehrer, P. M., Nolan, R. P., & Siepmann, M. (2023). Editorial: Integrated cardiovascular and neural system processes as potential mechanisms of behavior change. Frontiers in Psychiatry, 14, 1175691. https://doi.org/10.3389/fpsyt.2023.1175691

Berthoud, H.-R. (2008). The vagus nerve, food intake and obesity. Regulatory Peptides, 149(1–3), 15–25. https://doi.org/10.1016/j.regpep.2007.08.024

Berthoud, H.-R., Albaugh, V. L., & Neuhuber, W. L. (2021). Gut-brain communication and obesity: Understanding functions of the vagus nerve. Journal of Clinical Investigation, 131(10), e143770. https://doi.org/10.1172/JCI143770

Borges, U., Laborde, S., & Raab, M. (2019). Influence of transcutaneous vagus nerve stimulation on cardiac vagal activity: Not different from sham stimulation and no effect of stimulation intensity. PLOS ONE, 14(10), e0223848. https://doi.org/10.1371/journal.pone.0223848

Borgmann, D., Rigoux, L., Kuzmanovic, B., Edwin Thanarajah, S., Münte, T. F., Fenselau, H., & Tittgemeyer, M. (2021). Technical Note: Modulation of fMRI brainstem responses by transcutaneous vagus nerve stimulation. NeuroImage, 244, 118566. https://doi.org/10.1016/j.neuroimage.2021.118566

Bretherton, B., Atkinson, L., Murray, A., Clancy, J., Deuchars, S., & Deuchars, J. (2019). Effects of transcutaneous vagus nerve stimulation in individuals aged 55 years or above: Potential benefits of daily stimulation. Aging, 11(14), 4836–4857. https://doi.org/10.18632/aging.102074

Buchman, T. G., Stein, P. K., & Goldstein, B. (2002). Heart rate variability in critical illness and critical care: Current Opinion in Critical Care, 8(4), 311–315. https://doi.org/10.1097/00075198-200208000-00007

Burger, A. M., D’Agostini, M., Verkuil, B., & Van Diest, I. (2020). Moving beyond belief: A narrative review of potential biomarkers for transcutaneous vagus nerve stimulation. Psychophysiology, 57(6). https://doi.org/10.1111/psyp.13571

Burger, A. M., Van der Does, W., Thayer, J. F., Brosschot, J. F., & Verkuil, B. (2019). Transcutaneous vagus nerve stimulation reduces spontaneous but not induced negative thought intrusions in high worriers. Biological Psychology, 142, 80–89. https://doi.org/10.1016/j.biopsycho.2019.01.014

Butt, M. F., Albusoda, A., Farmer, A. D., & Aziz, Q. (2020). The anatomical basis for transcutaneous auricular vagus nerve stimulation. Journal of Anatomy, 236(4), 588– 611. https://doi.org/10.1111/joa.13122

Capilupi, M. J., Kerath, S. M., & Becker, L. B. (2020). Vagus Nerve Stimulation and the Cardiovascular System. Cold Spring Harbor Perspectives in Medicine, 10(2), a034173. https://doi.org/10.1101/cshperspect.a034173

Chen, M., Yu, L., Ouyang, F., Liu, Q., Wang, Z., Wang, S., Zhou, L., Jiang, H., & Zhou, S. (2015). The right side or left side of noninvasive transcutaneous vagus nerve stimulation: Based on conventional wisdom or scientific evidence? International Journal of Cardiology, 187, 44–45. https://doi.org/10.1016/j.ijcard.2015.03.351

Chen, Y., Lu, X., & Hu, L. (2023). Transcutaneous Auricular Vagus Nerve Stimulation Facilitates Cortical Arousal and Alertness. International Journal of Environmental Research and Public Health, 20(2), 1402. https://doi.org/10.3390/ijerph20021402

Clancy, J. A., Mary, D. A., Witte, K. K., Greenwood, J. P., Deuchars, S. A., & Deuchars, J. (2014). Non-invasive Vagus Nerve Stimulation in Healthy Humans Reduces Sympathetic Nerve Activity. Brain Stimulation, 7(6), 871–877. https://doi.org/10.1016/j.brs.2014.07.031

Coopmans, C., Zhou, T. L., Henry, R. M. A., Heijman, J., Schaper, N. C., Koster, A., Schram, M. T., van der Kallen, C. J. H., Wesselius, A., den Engelsman, R. J. A., Crijns, H. J. G. M., & Stehouwer, C. D. A. (2020). Both Prediabetes and Type 2 Diabetes Are Associated With Lower Heart Rate Variability: The Maastricht Study. Diabetes Care, 43(5), 1126–1133. https://doi.org/10.2337/dc19-2367

D’Agostini, M., Burger, A. M., Franssen, M., Perkovic, A., Claes, S., von Leupoldt, A., Murphy, P. R., & Van Diest, I. (2023). Short bursts of transcutaneous auricular vagus nerve stimulation enhance evoked pupil dilation as a function of stimulation parameters. Cortex, 159, 233–253. https://doi.org/10.1016/j.cortex.2022.11.012

De Couck, M., Cserjesi, R., Caers, R., Zijlstra, W. P., Widjaja, D., Wolf, N., Luminet, O., Ellrich, J., & Gidron, Y. (2017). Effects of short and prolonged transcutaneous vagus nerve stimulation on heart rate variability in healthy subjects. Autonomic Neuroscience, 203, 88–96. https://doi.org/10.1016/j.autneu.2016.11.003

Dickerson, S. S., & Kemeny, M. E. (2004). Acute Stressors and Cortisol Responses: A Theoretical Integration and Synthesis of Laboratory Research. Psychological Bulletin, 130(3), 355–391. https://doi.org/10.1037/0033-2909.130.3.355

Eckberg, D. L. (1983). Human sinus arrhythmia as an index of vagal cardiac outflow. Journal of Applied Physiology, 54(4), 961–966. https://doi.org/10.1152/jappl.1983.54.4.961

Farmer, A. D., Strzelczyk, A., Finisguerra, A., Gourine, A. V., Gharabaghi, A., Hasan, A., Burger, A. M., Jaramillo, A. M., Mertens, A., Majid, A., Verkuil, B., Badran, B. W., Ventura-Bort, C., Gaul, C., Beste, C., Warren, C. M., Quintana, D. S., Hämmerer, D., Freri, E., … Koenig, J. (2021). International Consensus Based Review and Recommendations for Minimum Reporting Standards in Research on Transcutaneous Vagus Nerve Stimulation (Version 2020). Frontiers in Human Neuroscience, 14, 568051. https://doi.org/10.3389/fnhum.2020.568051

Ferstl, M., Teckentrup, V., Lin, W. M., Kräutlein, F., Kühnel, A., Klaus, J., Walter, M., & Kroemer, N. B. (2021). Non-invasive vagus nerve stimulation boosts mood recovery after effort exertion. Psychological Medicine, 1–11. https://doi.org/10.1017/S0033291720005073

Forte, G., Favieri, F., Leemhuis, E., De Martino, M. L., Giannini, A. M., De Gennaro, L., Casagrande, M., & Pazzaglia, M. (2022). Ear your heart: Transcutaneous auricular vagus nerve stimulation on heart rate variability in healthy young participants. PeerJ, 10, e14447. https://doi.org/10.7717/peerj.14447

Frangos, E., Ellrich, J., & Komisaruk, B. R. (2015). Non-invasive Access to the Vagus Nerve Central Projections via Electrical Stimulation of the External Ear: FMRI Evidence in Humans. Brain Stimulation, 8(3), 624–636. https://doi.org/10.1016/j.brs.2014.11.018

Galli, R., Limbruno, U., Pizzanelli, C., Sean Giorgi, F., Lutzemberger, L., Strata, G., Pataleo, L., Mariani, M., Iudice, A., & Murri, L. (2003). Analysis of RR variability in drug-resistant epilepsy patients chronically treated with vagus nerve stimulation. Autonomic Neuroscience, 107(1), 52–59. https://doi.org/10.1016/S1566-0702(03)00081-X

Gauthey, A., Morra, S., van de Borne, P., Deriaz, D., Maes, N., & le Polain de Waroux, J.-B. (2020). Sympathetic Effect of Auricular Transcutaneous Vagus Nerve Stimulation on Healthy Subjects: A Crossover Controlled Clinical Trial Comparing Vagally Mediated and Active Control Stimulation Using Microneurography. Frontiers in Physiology, 11, 599896. https://doi.org/10.3389/fphys.2020.599896

Geng, D., Liu, X., Wang, Y., & Wang, J. (2022). The effect of transcutaneous auricular vagus nerve stimulation on HRV in healthy young people. PLOS ONE, 17(2), e0263833. https://doi.org/10.1371/journal.pone.0263833

Giraudier, M., Ventura-Bort, C., Burger, A. M., Claes, N., D’Agostini, M., Fischer, R., Franssen, M., Kaess, M., Koenig, J., Liepelt, R., Nieuwenhuis, S., Sommer, A., Usichenko, T., Van Diest, I., von Leupoldt, A., Warren, C. M., & Weymar, M. (2022). Evidence for a modulating effect of transcutaneous auricular vagus nerve stimulation (taVNS) on salivary alpha-amylase as indirect noradrenergic marker: A pooled mega-analysis. Brain Stimulation, 15(6), 1378–1388. https://doi.org/10.1016/j.brs.2022.09.009

Gourine, A. V., Machhada, A., Trapp, S., & Spyer, K. M. (2016). Cardiac vagal preganglionic neurones: An update. Autonomic Neuroscience, 199, 24–28. https://doi.org/10.1016/j.autneu.2016.06.003

Grossman, P., & Taylor, E. W. (2007). Toward understanding respiratory sinus arrhythmia: Relations to cardiac vagal tone, evolution and biobehavioral functions. Biological Psychology, 74(2), 263–285. https://doi.org/10.1016/j.biopsycho.2005.11.014

Havel, P. J. (2001). Peripheral Signals Conveying Metabolic Information to the Brain: Short-Term and Long-Term Regulation of Food Intake and Energy Homeostasis. Experimental Biology and Medicine, 226(11), 963–977. https://doi.org/10.1177/153537020122601102

Hedman, A. E., Hartikainen, J. E. K., Tahvanainen, K. U. O., & Hakumäki, M. O. K. (1995). The high frequency component of heart rate variability reflects cardiac parasympathetic modulation rather than parasympathetic ‘tone’. Acta Physiologica Scandinavica, 155(3), 267–273. https://doi.org/10.1111/j.1748-1716.1995.tb09973.x

Hong, G.-S., Pintea, B., Lingohr, P., Coch, C., Randau, T., Schaefer, N., Wehner, S., Kalff, J. C., & Pantelis, D. (2019). Effect of transcutaneous vagus nerve stimulation on muscle activity in the gastrointestinal tract (transVaGa): A prospective clinical trial. International Journal of Colorectal Disease, 34(3), 417–422. https://doi.org/10.1007/s00384-018-3204-6

Huang, J., Wang, Y., Jiang, D., Zhou, J., & Huang, X. (2010). The sympathetic-vagal balance against endotoxemia. Journal of Neural Transmission, 117(6), 729–735. https://doi.org/10.1007/s00702-010-0407-6

Jansen, K., Vandeput, S., Milosevic, M., Ceulemans, B., Van Huffel, S., Brown, L., Penders, J., & Lagae, L. (2011). Autonomic effects of refractory epilepsy on heart rate variability in children: Influence of intermittent vagus nerve stimulation: Heart Rate Variability in Refractory Epilepsy. Developmental Medicine & Child Neurology, 53(12), 1143–1149. https://doi.org/10.1111/j.1469-8749.2011.04103.x

Jensen, M. K., Andersen, S. S., Andersen, S. S., Liboriussen, C. H., Kristensen, S., & Jochumsen, M. (2022). Modulating Heart Rate Variability through Deep Breathing Exercises and Transcutaneous Auricular Vagus Nerve Stimulation: A Study in Healthy Participants and in Patients with Rheumatoid Arthritis or Systemic Lupus Erythematosus. Sensors, 22(20), 7884. https://doi.org/10.3390/s22207884

Keute, M., Machetanz, K., Berelidze, L., Guggenberger, R., & Gharabaghi, A. (2021). Neuro-cardiac coupling predicts transcutaneous auricular vagus nerve stimulation effects. Brain Stimulation, 14(2), 209–216. https://doi.org/10.1016/j.brs.2021.01.001

Kim, A. Y., Marduy, A., de Melo, P. S., Gianlorenco, A. C., Kim, C. K., Choi, H., Song, J.-J., & Fregni, F. (2022). Safety of transcutaneous auricular vagus nerve stimulation (taVNS): A systematic review and meta-analysis. Scientific Reports, 12(1), 22055. https://doi.org/10.1038/s41598-022-25864-1

Koenig, J., Parzer, P., Haigis, N., Liebemann, J., Jung, T., Resch, F., & Kaess, M. (2021). Effects of acute transcutaneous vagus nerve stimulation on emotion recognition in adolescent depression. Psychological Medicine, 51(3), 511–520. https://doi.org/10.1017/S0033291719003490

Kozorosky, E. M., Lee, C. H., Lee, J. G., Nunez Martinez, V., Padayachee, L. E., & Stauss, H. M. (2022). Transcutaneous auricular vagus nerve stimulation augments postprandial inhibition of ghrelin. Physiological Reports, 10(8). https://doi.org/10.14814/phy2.15253

Laborde, S., Mosley, E., & Thayer, J. F. (2017). Heart Rate Variability and Cardiac Vagal Tone in Psychophysiological Research – Recommendations for Experiment Planning, Data Analysis, and Data Reporting. Frontiers in Psychology, 08. https://doi.org/10.3389/fpsyg.2017.00213

Lee, S. W., Anderson, A., Guzman, P. A., Nakano, A., Tolkacheva, E. G., & Wickman, K. (2018). Atrial GIRK Channels Mediate the Effects of Vagus Nerve Stimulation on Heart Rate Dynamics and Arrhythmogenesis. Frontiers in Physiology, 9, 943. https://doi.org/10.3389/fphys.2018.00943

Lewine, J. D., Paulson, K., Bangera, N., & Simon, B. J. (2019). Exploration of the Impact of Brief Noninvasive Vagal Nerve Stimulation on EEG and Event-Related Potentials. Neuromodulation: Technology at the Neural Interface, 22(5), 564–572. https://doi.org/10.1111/ner.12864

Lloyd, B., Wurm, F., De Kleijn, R., & Nieuwenhuis, S. (2023). *Short-term transcutaneous vagus nerve stimulation increases pupil size but does not affect EEG alpha power: A replication* [Preprint]. Neuroscience. https://doi.org/10.1101/2023.03.08.531479

Lu, C. L., Zou, X., Orr, W. C., & Chen, J. D. (1999). Postprandial changes of sympathovagal balance measured by heart rate variability. Digestive Diseases and Sciences, 44(4), 857–861. https://doi.org/10.1023/a:1026698800742

Machetanz, K., Berelidze, L., Guggenberger, R., & Gharabaghi, A. (2021). Transcutaneous auricular vagus nerve stimulation and heart rate variability: Analysis of parameters and targets. Autonomic Neuroscience, 236, 102894. https://doi.org/10.1016/j.autneu.2021.102894

Maniscalco, J. W., & Rinaman, L. (2018). Vagal Interoceptive Modulation of Motivated Behavior. Physiology, 33(2), 151–167. https://doi.org/10.1152/physiol.00036.2017

Marmerstein, J. T., McCallum, G. A., & Durand, D. M. (2021). Direct measurement of vagal tone in rats does not show correlation to HRV. Scientific Reports, 11(1), 1210. https://doi.org/10.1038/s41598-020-79808-8

Neuser, M. P., Teckentrup, V., Kühnel, A., Hallschmid, M., Walter, M., & Kroemer, N. B. (2020). Vagus nerve stimulation boosts the drive to work for rewards. Nature Communications, 11(1), 3555. https://doi.org/10.1038/s41467-020-17344-9

Ng, G. A., Brack, K. E., & Coote, J. H. (2001). Effects of Direct Sympathetic and Vagus Nerve Stimulation on the Physiology of the Whole Heart—A Novel Model of Isolated Langendorff Perfused Rabbit Heart with Intact Dual Autonomic Innervation. Experimental Physiology, 86(3), 319–329. https://doi.org/10.1113/eph8602146

Ohara, K., Okita, Y., Kouda, K., Mase, T., Miyawaki, C., & Nakamura, H. (2015). Cardiovascular response to short-term fasting in menstrual phases in young women: An observational study. BMC Women’s Health, 15(1), 67. https://doi.org/10.1186/s12905-015-0224-z

Oostenveld, R., Fries, P., Maris, E., & Schoffelen, J.-M. (2011). FieldTrip: Open source software for advanced analysis of MEG, EEG, and invasive electrophysiological data. Computational Intelligence and Neuroscience, 2011, 156869. https://doi.org/10.1155/2011/156869

Pascual, F. T. (2015). Vagus nerve stimulation and late-onset bradycardia and asystole: Case report. Seizure, 26, 5–6. https://doi.org/10.1016/j.seizure.2015.01.006

Petzschner, F. H., Garfinkel, S. N., Paulus, M. P., Koch, C., & Khalsa, S. S. (2021). Computational Models of Interoception and Body Regulation. Trends in Neurosciences, 44(1), 63–76. https://doi.org/10.1016/j.tins.2020.09.012

Peuker, E. T., & Filler, T. J. (2002). The nerve supply of the human auricle. Clinical Anatomy, 15(1), 35–37. https://doi.org/10.1002/ca.1089

Pomeranz, B., Macaulay, R. J., Caudill, M. A., Kutz, I., Adam, D., Gordon, D., Kilborn, K. M., Barger, A. C., Shannon, D. C., Cohen, R. J., & Et, Al. (1985). Assessment of autonomic function in humans by heart rate spectral analysis. American Journal of Physiology-Heart and Circulatory Physiology, 248(1), H151–H153. https://doi.org/10.1152/ajpheart.1985.248.1.H151

Prescott, S. L., & Liberles, S. D. (2022). Internal senses of the vagus nerve. Neuron, 110(4), 579–599. https://doi.org/10.1016/j.neuron.2021.12.020

Quintana, D. S., & Williams, D. R. (2018). Bayesian alternatives for common null-hypothesis significance tests in psychiatry: A non-technical guide using JASP. BMC Psychiatry, 18(1), 178. https://doi.org/10.1186/s12888-018-1761-4

Rebollo, I., Devauchelle, A.-D., Béranger, B., & Tallon-Baudry, C. (2018). Stomach-brain synchrony reveals a novel, delayed-connectivity resting-state network in humans. ELife, 7, e33321. https://doi.org/10.7554/eLife.33321

Redgrave, J., Day, D., Leung, H., Laud, P. J., Ali, A., Lindert, R., & Majid, A. (2018). Safety and tolerability of Transcutaneous Vagus Nerve stimulation in humans; a systematic review. Brain Stimulation, 11(6), 1225–1238. https://doi.org/10.1016/j.brs.2018.08.010

Rong, P., Liu, A., Zhang, J., Wang, Y., He, W., Yang, A., Li, L., Ben, H., Li, L., Liu, H., Wu, P., Liu, R., Zhao, Y., Zhang, J., Huang, F., Li, X., & Zhu, B. (2014). Transcutaneous vagus nerve stimulation for refractory epilepsy: A randomized controlled trial. Clinical Science, CS20130518. https://doi.org/10.1042/CS20130518

Sclocco, R., Garcia, R. G., Kettner, N. W., Isenburg, K., Fisher, H. P., Hubbard, C. S., Ay, I., Polimeni, J. R., Goldstein, J., Makris, N., Toschi, N., Barbieri, R., & Napadow, V. (2019). The influence of respiration on brainstem and cardiovagal response to auricular vagus nerve stimulation: A multimodal ultrahigh-field (7T) fMRI study. Brain Stimulation, 12(4), 911–921. https://doi.org/10.1016/j.brs.2019.02.003

Shaffer, F., & Ginsberg, J. P. (2017). An Overview of Heart Rate Variability Metrics and Norms. Frontiers in Public Health, 5, 258. https://doi.org/10.3389/fpubh.2017.00258

Shankar, R., Olotu, V. O., Cole, N., Sullivan, H., & Jory, C. (2013). Case report: Vagal nerve stimulation and late onset asystole. Seizure, 22(4), 312–314. https://doi.org/10.1016/j.seizure.2012.12.011

Sharon, O., Fahoum, F., & Nir, Y. (2021). Transcutaneous Vagus Nerve Stimulation in Humans Induces Pupil Dilation and Attenuates Alpha Oscillations. The Journal of Neuroscience, 41(2), 320–330. https://doi.org/10.1523/JNEUROSCI.1361-20.2020

Šinkovec, M., Trobec, R., Kamenski, T., Jerman, N., & Meglič, B. (2023). Hemodynamic responses to low-level transcutaneous auricular nerve stimulation in young volunteers. IBRO Neuroscience Reports, S2667242123000106. https://doi.org/10.1016/j.ibneur.2023.01.010

Soer, R., Six Dijkstra, M. W. M. C., Bieleman, H. J., Oosterveld, F. G. J., & Rijken, N. H. M. (2021). Influence of respiration frequency on heart rate variability parameters: A randomized cross-sectional study. Journal of Back and Musculoskeletal Rehabilitation, 34(6), 1063–1068. https://doi.org/10.3233/BMR-200190

Szulczewski, M. T. (2022). Transcutaneous Auricular Vagus Nerve Stimulation Combined With Slow Breathing: Speculations on Potential Applications and Technical Considerations. Neuromodulation: Technology at the Neural Interface, 25(3), 380–394. https://doi.org/10.1111/ner.13458

Teckentrup, V., Krylova, M., Jamalabadi, H., Neubert, S., Neuser, M. P., Hartig, R., Fallgatter, A. J., Walter, M., & Kroemer, N. B. (2021). Brain signaling dynamics after vagus nerve stimulation. NeuroImage, 245, 118679. https://doi.org/10.1016/j.neuroimage.2021.118679

Teckentrup, V., Neubert, S., Santiago, J. C. P., Hallschmid, M., Walter, M., & Kroemer, N. B. (2020). Non-invasive stimulation of vagal afferents reduces gastric frequency. Brain Stimulation, 13(2), 470–473. https://doi.org/10.1016/j.brs.2019.12.018

Thayer, J. F., Hansen, A. L., Saus-Rose, E., & Johnsen, B. H. (2009). Heart Rate Variability, Prefrontal Neural Function, and Cognitive Performance: The Neurovisceral Integration Perspective on Self-regulation, Adaptation, and Health. Annals of Behavioral Medicine, 37(2), 141–153. https://doi.org/10.1007/s12160-009-9101-z

Thayer, J. F., & Lane, R. D. (2007). The role of vagal function in the risk for cardiovascular disease and mortality. Biological Psychology, 74(2), 224–242. https://doi.org/10.1016/j.biopsycho.2005.11.013

Vest, A. N., Da Poian, G., Li, Q., Liu, C., Nemati, S., Shah, A. J., & Clifford, G. D. (2018). An open source benchmarked toolbox for cardiovascular waveform and interval analysis. Physiological Measurement, 39(10), 105004. https://doi.org/10.1088/1361-6579/aae021

Vosseler, A., Zhao, D., Fritsche, L., Lehmann, R., Kantartzis, K., Small, D. M., Peter, A., Häring, H.-U., Birkenfeld, A. L., Fritsche, A., Wagner, R., Preißl, H., Kullmann, S., & Heni, M. (2020). No modulation of postprandial metabolism by transcutaneous auricular vagus nerve stimulation: A cross-over study in 15 healthy men. Scientific Reports, 10(1), 20466. https://doi.org/10.1038/s41598-020-77430-2

Weise, D., Adamidis, M., Pizzolato, F., Rumpf, J.-J., Fricke, C., & Classen, J. (2015). Assessment of Brainstem Function with Auricular Branch of Vagus Nerve Stimulation in Parkinson’s Disease. PLOS ONE, 10(4), e0120786. https://doi.org/10.1371/journal.pone.0120786

Wolf, V., Kühnel, A., Teckentrup, V., Koenig, J., & Kroemer, N. B. (2021). Does transcutaneous auricular vagus nerve stimulation affect vagally mediated heart rate variability? A living and interactive Bayesian meta-analysis. Psychophysiology, 58(11). https://doi.org/10.1111/psyp.13933

Yakunina, N., Kim, S. S., & Nam, E.-C. (2017). Optimization of Transcutaneous Vagus Nerve Stimulation Using Functional MRI. Neuromodulation: Technology at the Neural Interface, 20(3), 290–300. https://doi.org/10.1111/ner.12541

Yoshida, K., Saku, K., Kamada, K., Abe, K., Tanaka-Ishikawa, M., Tohyama, T., Nishikawa, T., Kishi, T., Sunagawa, K., & Tsutsui, H. (2018). Electrical Vagal Nerve Stimulation Ameliorates Pulmonary Vascular Remodeling and Improves Survival in Rats With Severe Pulmonary Arterial Hypertension. JACC: Basic to Translational Science, 3(5), 657–671. https://doi.org/10.1016/j.jacbts.2018.07.007

