## Supplementary Information for "Non-invasive vagus nerve stimulation decreases vagally mediated heart rate variability"

### Supplementary Material A: The Milkshake recipe

The milkshake (basic recipe see Sun et al., 2015) contained 250g milk (3.5% fat, heat-treated; Gut & Günstig, Edeka, Germany), 50g cream (30% fat, heat-treated; Gut & Günstig, Edeka, Germany) and 50g syrup (Hershey's Syrup, The Hershey Company, USA). This corresponds to a nutrient composition of 9.4g protein (12%), 23.7g fat (31%), and 43.6g carbohydrates (57%). The milkshake has an energy of 400 kcal (1674.7 kJ) and a volume of 350 ml. To offer participants a milkshake they would like to consume, we offered three flavors (strawberry, chocolate, caramel) with comparable nutrient composition. The chosen flavor was used for all sessions throughout the experiment.

### Supplementary Material B:

Table S1. Mean und standard deviation of the applied stimulation intensities

| **Intensity in mA (mean, sd)** | **sham*** | **taVNS*** |
| --- | --- | --- |
| **Left** | 2.84 ± 1.09 | 2.40 ± 1.13 |
| **Right** | 2.95 ± 1.03 | 2.24 ± 1.00 |

*Note. **The stimulation intensities were matched between taVNS and sham on the perceptual level

### Supplementary Material C: Description for automatic artefact detection of the R peaks and R-R intervals

We evaluated whether the minimum distance between valid R peaks is at least 0.6 s (corresponding to a minimum HR of 36 bpm), and the minimum height of the peaks should be the median of the ECG energy profile (Lanata et al., 2015). Invalid peaks are identified by more than four times the median of the absolute difference between the amplitude of peaks. Invalid R-R intervals deviate by a factor of 0.6 from the mode of R-R intervals or by twice the mode of the R-R intervals. Finally, a Hampel filter, recommended as a moving window for nonlinear data cleaning (Ghaleb et al., 2018), was applied to search for deviations between consecutive intervals as an alternative measure of the distance from the median of its neighboring observations. All detected deviations and their consecutive R-R intervals were deleted from the signal. On average, 97.1% (1769s, range: 81% - 100%) of a 30 min block of ECG recordings was further analyzed to derive cardiovascular indices (except for one blocks with 956 s of ECG data). Overall, we carefully processed the ECG data and checked for artifacts to ensure the validity and accuracy of our results.

### Supplementary Material D: raw data of the HRV Addition (related to Figure 3)

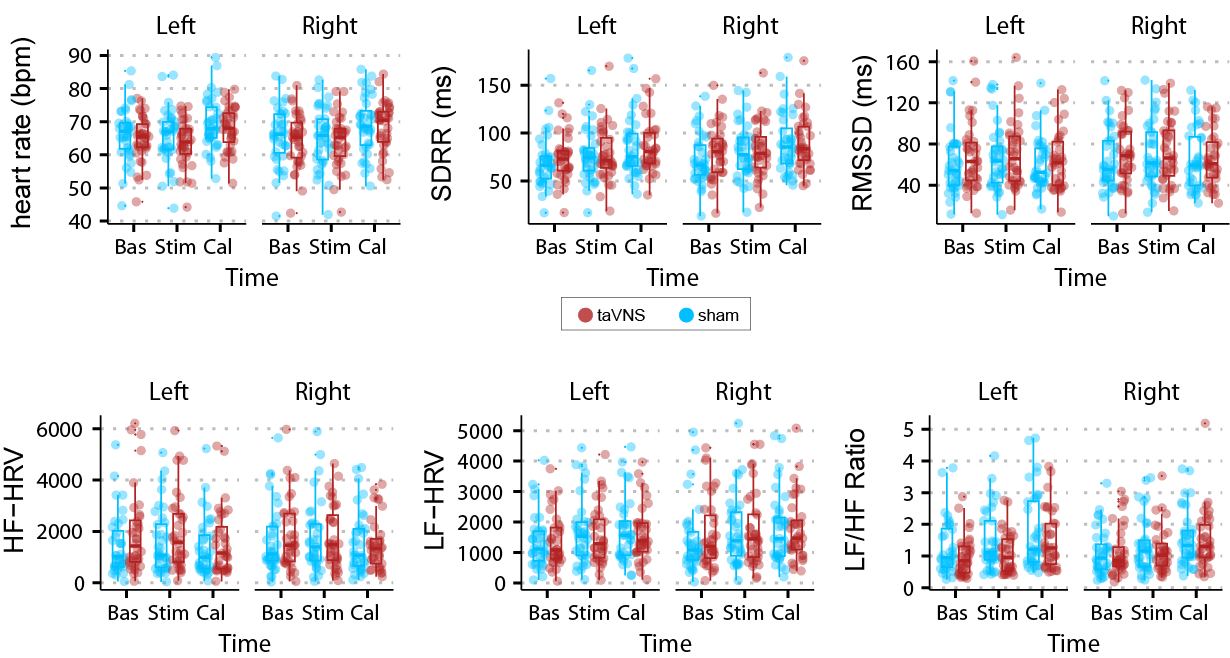

**Figure S1. taVNS decreases HRV.** Displayed are the raw data for HR and the different HRV indices over time (baseline vs. stimulation vs. caloric load), respectively, for the two stimulation sides (left vs. right) and the two stimulation conditions (taVNS in red vs. sham in blue). The dots show the average score of each participant, and the boxplots show the interquartile range (IQR), where the line inside the box is the median, and the whiskers represent the minimum and maximum values within 1.5 times the IQR.

### Supplementary Material E: Table for the results in figure 4 C-E

Table S2. Correlations between right and left side for taVNS or sham

| **Variable** | **Stimulation** | **R** | ***p*-value** |  | **R** | ***p*-value** |
| --- | --- | --- | --- | --- | --- | --- |
|  | | Stimulation | |  | Caloric Load | |
| HR | Sham | **0.37** | **.027** |  | 0.18 | .280 |
|  | taVNS | **0.43** | **.008** |  | 0.32 | .058 |
| RMSSD | Sham | 0.28 | .102 |  | 0.11 | .517 |
|  | taVNS | **0.42** | **.011** |  | **0.48** | **.003** |
| SDRR | Sham | 0.25 | .149 |  | **0.56** | **< .001** |
|  | taVNS | **0.39** | **.019** |  | **0.58** | **< .001** |
| HF HRV | Sham | 0.09 | .611 |  | -0.07 | .702 |
|  | taVNS | **0.43** | **.009** |  | **0.58** | **< .001** |
| LF/HF ratio | Sham | 0.32 | .053 |  | **0.46** | **.005** |
|  | taVNS | **0.49** | **.002** |  | **0.47** | **.004** |

*Note.* R = Pearson-Correlation-Coefficient

### Supplementary Material F:

Table S3. Results of difference in taVNS-induced effects between both phases

| **Variable** | **Contrast** | **mean** | **[CI_L_ , CI_U_]** |  | ***p*-value** |
| --- | --- | --- | --- | --- | --- |
| RMSSD | Stim vs.Cal | -1.04 | -4.26, -2.22 |  | .531 |
| SDRR | Stim vs.Cal | -0.19 | -3.63, -3.07 |  | .922 |
| HF HRV | Stim vs.Cal | -95.88 | -237.32, 41.26 |  | .171 |
| LF/HF ratio | Stim vs.Cal | 0.10 | -0.04, 0.23 |  | .152 |
| Heart rate | Stim vs.Cal | 0.60 | -0.03, 1.22 |  | .063 |

*Note.* CI_L_ = 95% lower bound of confidence Interval; CI_U_ = 95% upper bound of confidence Interval, *p* = p-value
